# Novel transcriptional activity and extensive allelic imbalance in the human MHC region

**DOI:** 10.1101/186809

**Authors:** Elizabeth Gensterblum, Weisheng Wu, Amr H Sawalha

## Abstract

The major histocompatibility complex (MHC) region encodes human leukocyte antigen (HLA) genes and is the most complex region in the human genome. The extensive polymorphic nature of the HLA hinders accurate localization and functional assessment of disease risk loci within this region. Using targeted capture sequencing and constructing individualized genomes for transcriptome alignment, we identified 908 novel transcripts within the human MHC region. These include 593 novel isoforms of known genes, 137 antisense strand RNAs, 119 novel long intergenic noncoding RNAs, and 5 transcripts of 3 novel putative protein coding human endogenous retrovirus genes. We revealed extensive allele-dependent expression imbalance involving 88% of all heterozygous transcribed single nucleotide polymorphisms throughout the MHC transcriptome. Further, we showed that the transcriptome within the MHC region can be defined by 14 distinct co-expression clusters, with evidence of co-regulation by unique transcription factors in at least 9 of these clusters. Our data suggest a very complex regulatory map of the human MHC, and can help uncover functional consequences of disease risk loci in this region.

## Introduction

The human major histocompatibility complex (MHC) is a highly complex polymorphic genomic region containing many important immune-related genes. This region includes the human leukocyte antigen (HLA) genes involved in both self-tolerance and antigen presentation. Polymorphisms within HLA genes have been associated with over 100 autoimmune diseases and cancers, and allelotyping of translated genes has been the focus of extensive research ^1–3^. Intergenic variants within the MHC region, which may serve a role in gene regulation, have also been associated with several immune-related diseases ^4, 5^. However, the role of these intergenic variants is often not clear because regulation within the MHC is incompletely understood. The MHC contains a complex regulatory network including *cis*-acting and *trans*-acting regulation bridging inside and outside the MHC region ^6, 7^. Due to both the high rate of polymorphism and the complex regulatory networks within the MHC, the functional effects of specific disease-associated variants are difficult to elucidate.

Long intergenic noncoding RNAs (lincRNAs) have been extensively implicated in transcriptional regulation by recruitment of regulatory proteins. These proteins proceed to regulate gene expression by epigenetic modification, such as DNA methylation and chromatin modification ^8, 9^. Recruitment of transcription factors by lincRNAs has also been described ^10^. However, many lincRNAs are weakly expressed, and therefore may not be detected by RNA sequencing that spans the entire transcriptome.

Sequence-specific enrichment by magnetic bead pulldown has previously been used to sequence HLA genes for allelotyping, and to elucidate regulatory regions of individual genes ^2, 4^. We performed targeted enrichment of the entire MHC region in primary human monocytes using sequence-specific capture probes, followed by high throughput sequencing of DNA and RNA (CaptureSeq) ^11^, to allow for deep sequencing coverage of the MHC region. We targeted the entire MHC, including both intergenic regions and known splice variants. We identified genetic variants, then constructed personalized genomes to accurately align RNA sequences. After enrichment and alignment to personalized genomes, we were able to detect lowly expressed transcripts, and by including all genomic regions, we were able to identify novel intergenic transcripts. We also comprehensively assessed allelic expression imbalance and revealed extensive allele-specific expression throughout the entire MHC, indicating that polymorphism is a mechanism of complex transcriptional regulation in this region.

## Results

We performed deep targeted genome and transcriptome sequencing of the human MHC region (Chromosome 6 (hg19): 28.5 Mb-33.5 Mb) in primary human monocytes. Constructing individualized genomes for aligning RNA sequencing reads generated by deep targeted transcriptome sequencing improved transcript alignment and characterization in is this complex polymorphic region. DNA sequence reads aligned against the reference genome human MHC region with a mean read depth in the tiled regions of 334.8 ± 84.3 in all samples (Figure S1). Genetic polymorphisms in each sample were identified and an individualized MHC genome in each sample was constructed. A total of 65,289 genetic variants relative to the reference genome were identified, including 62,449 genetic variants that are heterozygous in at least one sample.

Targeted RNA sequencing was performed following ribosomal RNA depletion, allowing for high density coverage with an average 36.3 million alignments to the MHC region. RNA sequence alignment was performed in an individualized SNP-tolerant mode using DNA sequencing data from each sample to allow alignment to polymorphic loci identified in each corresponding sample. This strategy significantly enhanced successful alignment of transcript reads to the polymorphic MHC region, which coupled with highly dense targeted RNA sequencing, allowed for accurate identification of known and novel transcripts in the MHC region, including transcripts with low expression levels.

We identified a total of 3,072 transcripts aligned to the human MHC region in human primary monocytes. Of these, 908 transcripts were identified as novel transcripts that were present in at least 2 independent samples (**Table S1**). This includes 517 and 76 novel coding and noncoding transcript isoforms of known genes, respectively. In addition, we identified 137 novel anti-sense strand transcripts, 119 lincRNA transcripts, 54 intronic noncoding RNA transcripts, and 5 transcripts of three novel coding genes (Figure 1).

**Figure 1:**
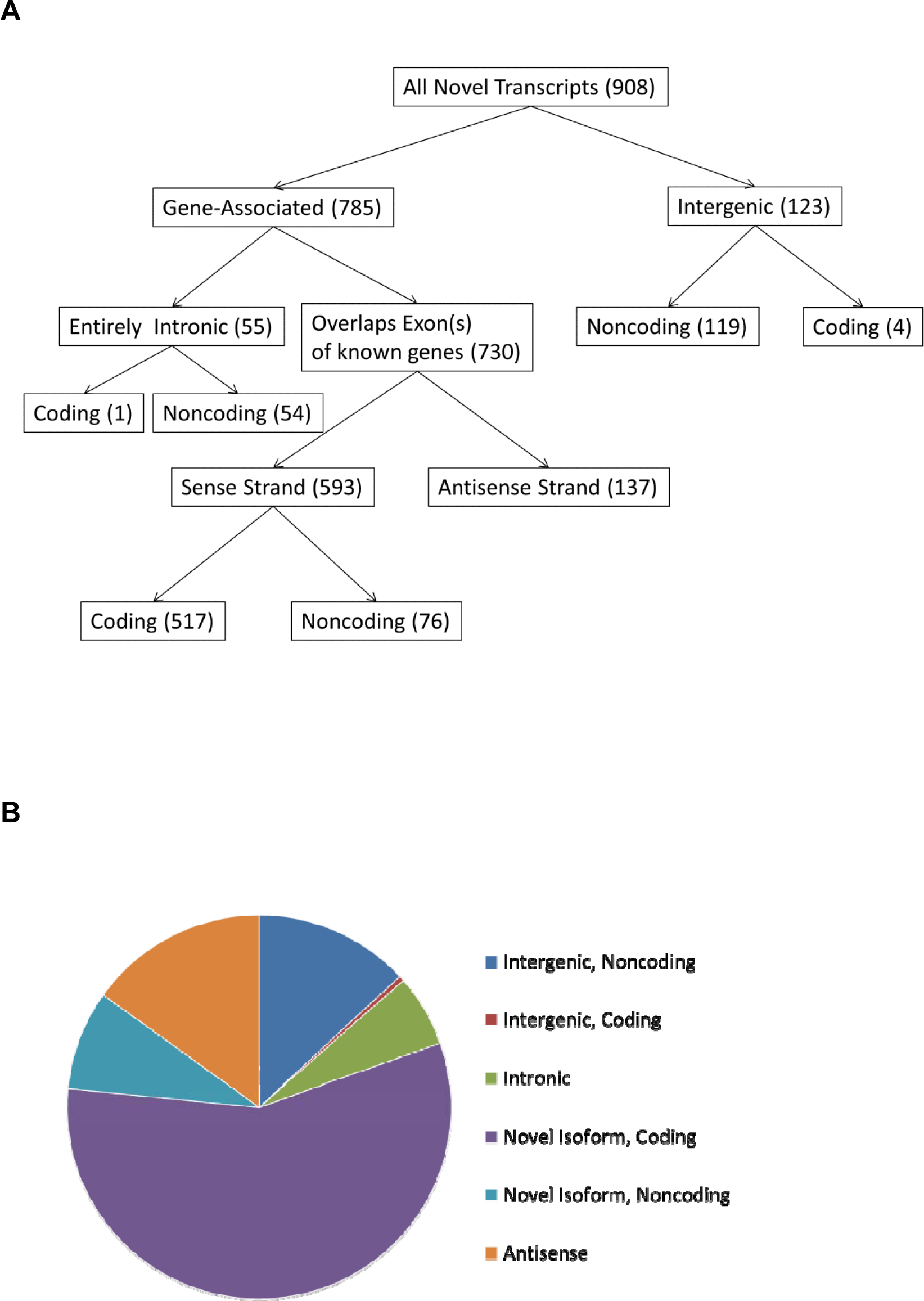
A flow chart (**A**) and pie chart (**B**) depicting and summarizing the filtering categories used to classify novel transcripts identified in this study. The categories for intronic, novel isoforms, antisense, and intergenic transcripts were defined via a CuffCompare annotation using the Comprehensive Gencode Resease 25 annotation (hg19) reference transcriptome. Coding potential of novel transcripts was predicted using CPAT.

Next, we evaluated the extent of allele-specific expression imbalance in MHC region transcripts that overlap with heterozygous single nucleotide polymorphisms identified using DNA sequencing. We show that 88% of heterozygous transcribed SNPs within the MHC region are associated with significant allele-dependent transcriptional imbalance, with 43% demonstrating extreme allele-dependent expression (>95% expression on either the reference or alternative allele) (Figure 2A). Indeed, allelic imbalance is observed in over 69% of all heterozygous SNPs identified in our study within the MHC region (**Table S2 and S3**). This remarkably extensive allele-specific expression pattern was non-stochastic and consistent across independent samples in heterozygous SNPs that are present in two or more samples (Figure 2B and 2C). While the overall number of heterozygous SNPs with evidence of allelic expression imbalance was highest in the HLA class II gene region within the MHC, the frequency of transcribed SNPs with allelic imbalance relative to all transcribed SNPs was consistent throughout the HLA regions within the MHC (Figure 3). To demonstrate allelic imbalance in a disease relevant locus in the MHC region, we examined the expression of novel transcripts that overlap with and include the SNP rs76546355 (rs116799036) localized between *HLA-B* and *MICA*. This genetic variant tags the most robust genetic association in Behçet’s disease ^5^. We show that rs76546355, previously thought to be intergenic, is expressed within four lincRNA transcripts localized between *HLA-B* and *MICA*. Importantly, these four transcripts are exclusively expressed from the DNA strand with the disease protective allele in this SNP. There was no expression of these transcripts from the disease risk allele strand in heterozygous individuals (Figure S3).

**Figure 2:**
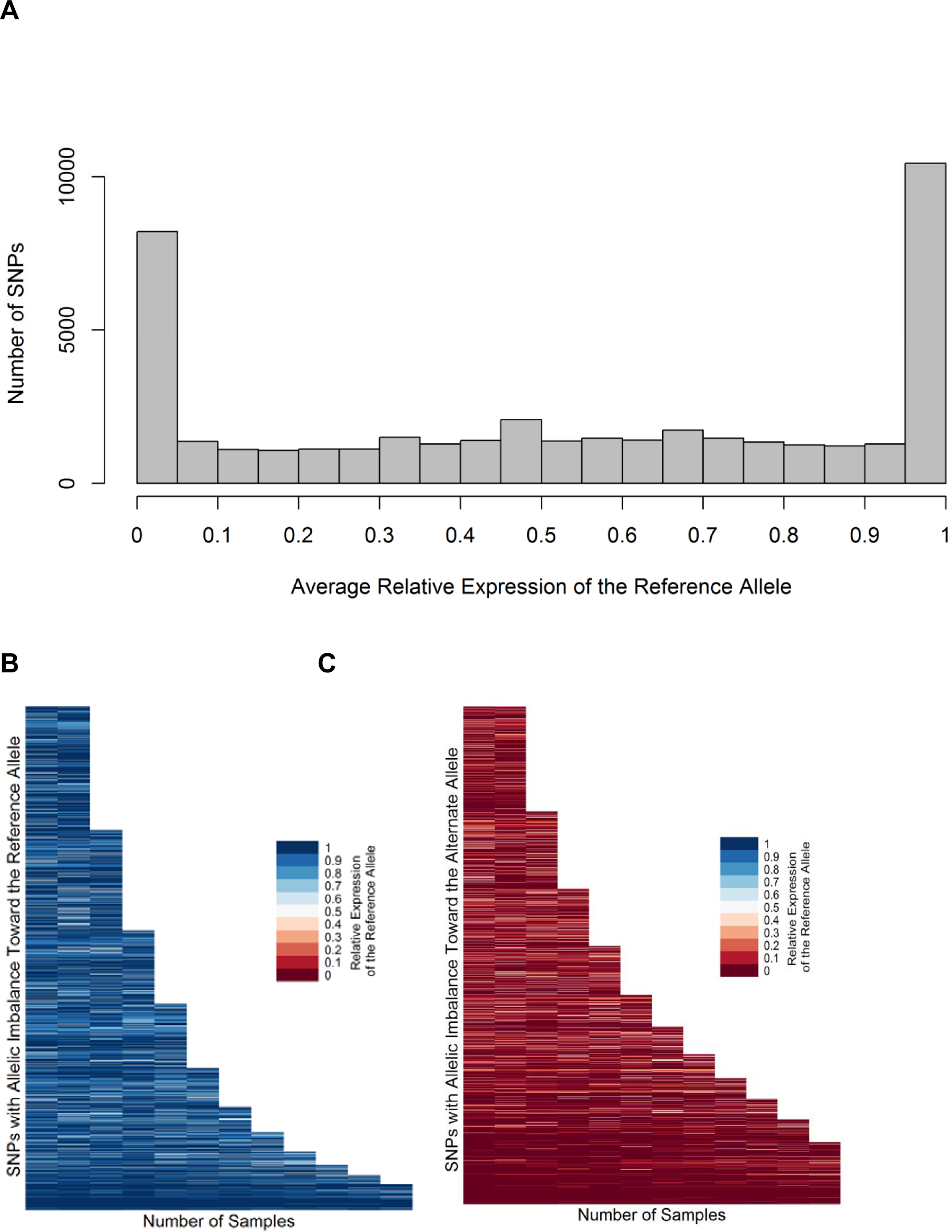
(**A**) Frequency distribution histogram of instances of allele-specific expression. The average relative expression (proportion of reads containing the reference allele) was calculated for all transcribed heterozygous SNPs identified in our study. Each bin spans a relative expression range of 0.05. (**B**) Variants in which the average relative expression of the reference allele is greater than 0.5, and in which the average relative expression is less than 0.5 (**C**). The relative expression of the reference allele in each SNP with allelic imbalance (binomial p<0.05) in two or more samples is represented on the Y axis. The reference allele is defined by the genotype of the reference genome, which is consistent across all samples. Relative expression ranges from 0 (red) to 1 (blue). The allelic imbalance of specific SNPs is shown to be highly consistent across individuals.

**Figure 3:**
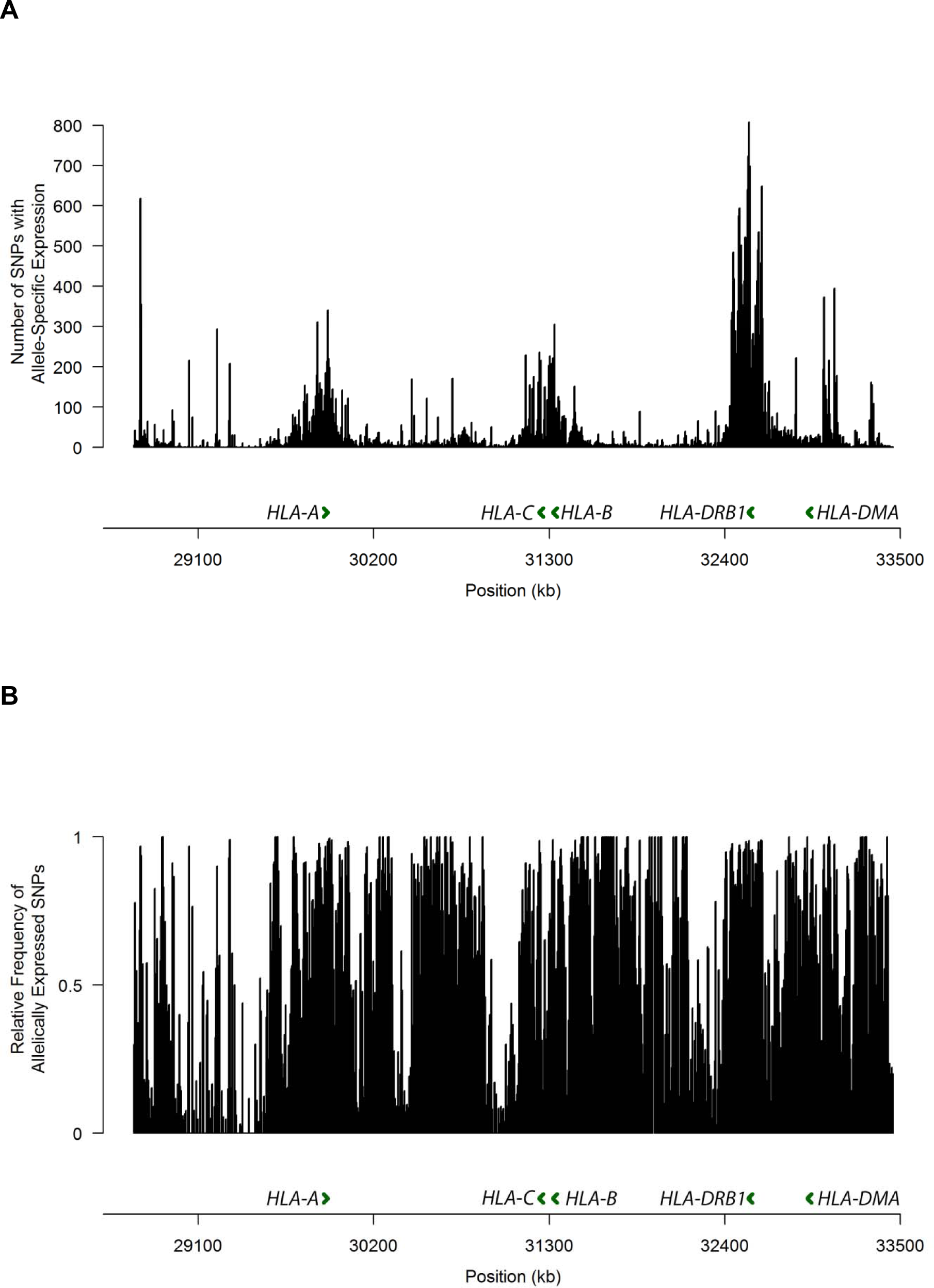
Histogram depicting the number of SNPs with allelic expression imbalance (**A**), and frequency of SNPs with allelic expression imbalance relative to all heterozygous SNPs detected in the MHC region (**B**). Each bin spans 5,000 base pairs.

We characterized the expression patterns of the transcripts within the MHC using a co-expression network analysis. We defined a co-expression network including all aligned transcripts, based on the normalized read counts across all twelve samples, using a weighted correlation network analysis (WGCNA)^12^. Based on this network, the transcripts were grouped into fourteen co-expression clusters, which do not localize to specific genomic regions within the MHC (Figure 4). Nevertheless, co-expression remains highly aggregated within individual clusters, and there is a high degree of separation between each cluster within the network (Figure S5).

**Figure 4:**
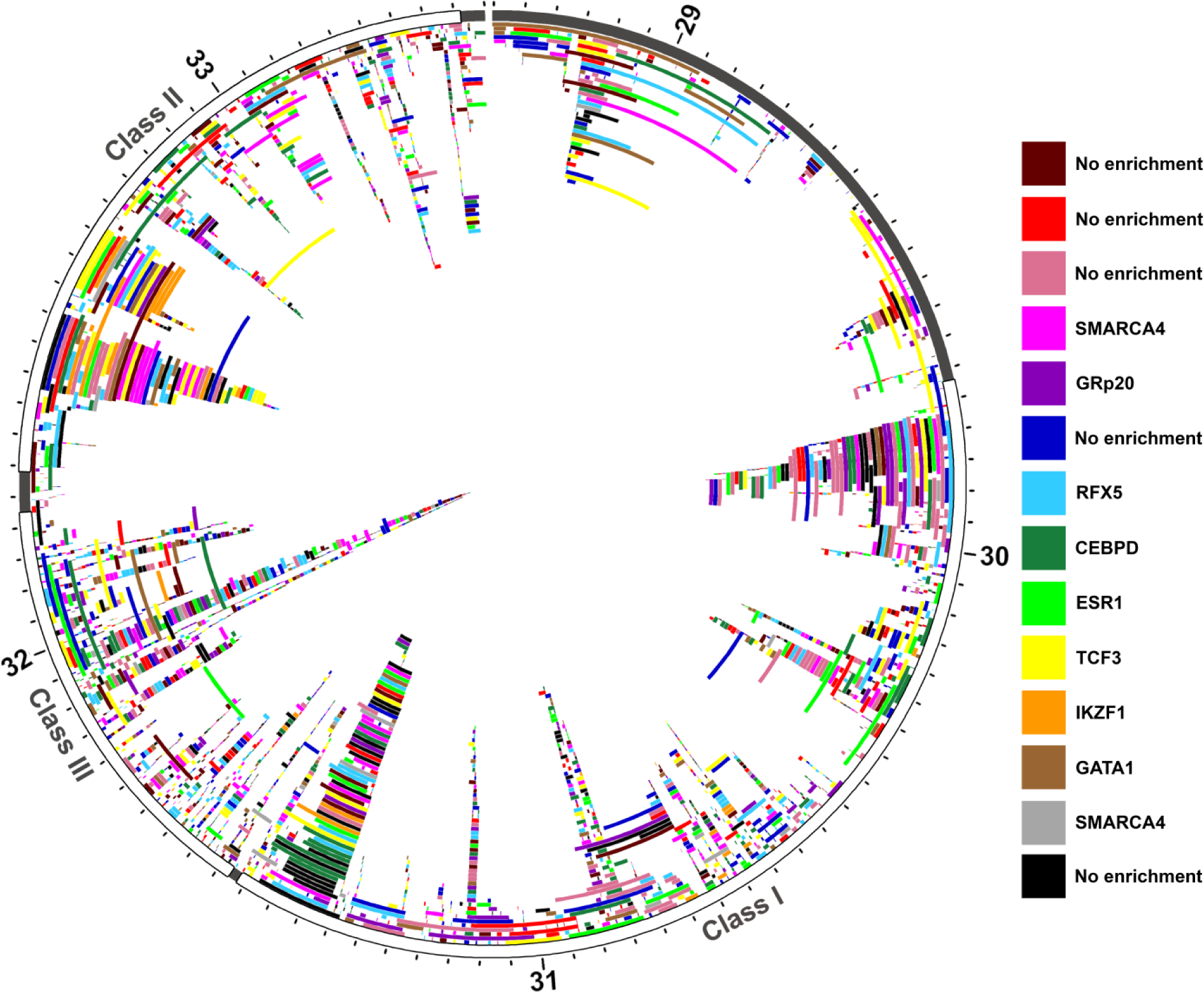
All unique transcripts plotted according to genomic position within the MHC region (chr6:28.7Mb-33.5Mb (hg19)). Chromosome six position (labelled in Mb) is plotted on the outer ideogram, and each MHC class is marked by color. The MHC class I is shown in red, MHC class II is green, and MHC class III is blue. Each aligned transcript, including novel transcripts, were grouped into co-expression clusters using the normalized read counts from each sequenced individual (n=12). Every transcript is plotted according to position, and colored according to cluster identity (red, dark red, orange, yellow, lime green, green, light blue, dark blue, purple, magenta, pink, black, grey, and brown). Multiple isoforms of the same gene can be found in the same co-expression cluster, but this is not a requirement and is never the case across all isoforms of a gene. There is no localization of transcripts within individual co-expression clusters based on genomic position. Nine of the fourteen clusters, however, were enriched for specific transcription factors, the most significantly enriched transcription factor for each cluster is listed.

We further described transcription factor binding site enrichment in each cluster. Of the fourteen co-expression clusters, nine were enriched for specific transcription factors (OR>1, p<0.05). For these clusters, the transcription factor binding sites most significantly enriched were TCF3 (OR: 2.17, p: 8.06 x 10^−7^), ESR1 (OR: 2.72, p: 2.21 x 10^−5^), RFX5 (OR: 1.68, p: 4.27 x 10^−5^), SMARCA4 (OR: 2.56, p: 2.10 x 10^−4^), GATA1 (OR:2.08, p: 2.25 x 10^−4^), IKZF1 (OR: 4.80, p: 6.92 x 10^−4^), GRp20 (OR: 2.57, p: 2.39 x 10^−3^), CEBPD (OR: 1.52, p: 5.44 x 10^−3^), and SMARCA4 (OR: 3.68, p: 6.74 x 10^−3^), respectively. The enrichment of these specific transcription factor binding sites suggests that these nine clusters may show co-expression due to transcription factor-dependent co-regulation.

We identified 3 novel genes with an open reading frame that are predicted to be protein coding within the human MHC region, and demonstrate gene expression at the mRNA level. Using protein function and structure prediction algorithms, two of the three coding genes we identified are predicted with moderate and very high certainty to be novel endogenous retroviral *pol* and *gag* genes, respectively (Figure S4). The structure and function of the third gene could not be predicted. The predicted amino acid sequences of all three genes were aligned to the human proteome using protein-protein Blast^13^. Based on the homology between each novel sequence and the HERVs to which it is aligned, we predict that these genes are retroviral *pol*, *gag*, and *gag* proteins, respectively (**Table S5)**.

## Discussion

Variation within the MHC contributes to genetic risk of immune and inflammatory disease. However, this region is characterized by complex variation patterns that complicate identifying causal variants and their direct effects on disease etiology ^14^. Moreover, these complex variation patterns play a role in the complex alternative splicing and gene regulation networks that have been described in this region ^7, 15^. Quantification of MHC transcription by RNA sequencing has been limited by both the high rate of polymorphism and the high rate of splice variants, resulting in limitations in RNA sequence alignment ^16^. Using individualized genomes to map RNA sequencing reads, we accurately measured gene and splice variant expression within the MHC, which can be used to further elucidate the functional effects of variations relevant to disease.

Sequence variation can affect the expression of transcripts by interfering with *cis*-regulation, such as altering promoter or enhancer activity, altering DNA methylation patterns, or altering the sequence of regulatory RNAs. Variants linked to these *cis*-effects (*cis*-eQTL) affect expression in an allele-specific manner. Haplotype-specific gene expression within the HLA, and allelic imbalance linked to *cis* eQTLs in autoimmunity have been previously described ^4, 17^. Our findings suggest extensive allele-specific expression throughout the MHC, which involves 88% of all transcribed SNPs in this region.

Many lincRNA transcripts are expressed at low levels, rendering them undetectable without sequence enrichment. By targeting the MHC region using sequence-specific capture probes, we identified novel noncoding transcripts throughout the region. As lincRNAs have been implicated in transcriptional regulation, this suggests a far more complex regulatory network within the MHC than what has been previously described. Variation within the MHC further affects the patterns of transcription regulation, due to allelic imbalance as we demonstrate.

When we compare the expression patterns of all transcripts across all twelve sequenced individuals, a pattern of co-expression develops. While the co-expression of genes does not intrinsically imply co-regulation, regulation by the same transcription factors is one mechanism by which co-expression can occur. After quantifying the enrichment of the transcription factors binding to the promoter regions of the transcripts in each cluster, we found that nine of the fourteen clusters were enriched for specific transcription factors. This suggests that regulation by these transcription factors may play a role in the expression patterns of the transcripts in each cluster. Some of the enriched transcription factors identified play a role in specific immunological processes. For example, one of these co-expression clusters was found to be enriched for RFX5, a transcription factor that activates MHC class II expression by enhancing CIITA activity^18^. Another transcription factor, enriched in a different cluster, CEBPD is directly involved in promoting macrophage activation, M1 macrophage polarization, and pro-inflammatory cytokine production in macrophages^19^. The transcription factor GATA1 is involved in dendritic cell differentiation and survival^20^. These enriched transcription factors each have a unique role in monocyte differentiation, suggesting that they have a role not only in determining co-expression patterns of transcripts, but in downstream determination of cellular phenotypes.

We found five novel putative coding transcripts, identifying three novel human endogenous retroviral elements (ERVs). ERVs comprise 8% of the human genome^21^. Though mutations have silenced the expression of the majority of these elements, approximately 7% of all known ERVs are transcriptionally active^22^. Moreover, mutations in these elements have been linked to diseases, including cancer^15^ and multiple sclerosis ^23^. Translated ERVs have been shown to play a role in lymphocyte activation, and transcribed ERVs play a role in transcriptional regulation ^24^. The exact function of these novel ERVs, and their precise effects on transcription and immune function, has yet to be fully elucidated.

In summary, we performed deep sequencing of both the genome and the transcriptome, targeting the MHC region with sequence-specific capture probes in human monocytes. We accurately identified and quantified the expression of 908 novel transcripts in this region, including 123 transcripts aligning to regions previously thought to be intergenic. In addition, we uncovered extensive allele-specific expression imbalance within the MHC region, which appears to be non-stochastic, suggesting complex *cis*-acting transcriptional regulation throughout the human MHC. This allelic imbalance can have functional consequences upon disease risk loci within the MHC region.

## Materials and Methods

### Probe Design

Sequence-specific capture probes were designed to target the complete reference genomic sequence of the MHC region (chr6:28.5 Mb-33.5 Mb, hg19), as well as splice sites for known transcripts the region contains. By including intergenic and intronic genomic regions, sequences that overlap with previously unannotated regions could be captured and subsequently sequenced; moreover, the same set of probes were designed to enable us to capture both DNA and RNA. This pool of 75 base long capture probes was designed to target 35,895 sequences throughout the region. For the main reference allele, probes directly overlapped 75.9% of the genome, with 88% estimated total sequence coverage. However, because the MHC region is highly polymorphic, the seven alternative reference haplotypes for the MHC were used in addition to the reference allele to design probes targeting all reference genomic sequences in this region. In total, this region, including all alternative haplotypes, was 65.4% covered by the probes, and had a 75.7% estimated net coverage. Of the total target region, including alternate haplotypes, 10% was not covered due to shared homology with other parts of the genome, while 14.2% was not covered because of incomplete sequence information in the alternative haplotypes.

### Isolation of Primary Monocyte DNA and RNA

Peripheral blood mononuclear cells (PBMCs) isolated from 12 healthy individuals were initially collected by density gradient centrifugation and immediately stored in liquid nitrogen. Cells were thawed, treated with 25 U/mL benzonase, and incubated at 37°C in RPMI/10% heat inactivated fetal bovine serum for 90 minutes. Thawed PBMCs had a minimum viability of 90%, with an average viability of 98.1% ± 3.6%, measured by tryphan blue staining. Primary monocytes were then isolated from thawed peripheral blood mononuclear cells via negative selection using the Pan Monocyte Isolation Kit, following the manufacturer’s instructions (Miltenyi Biotec Inc., San Diego California). The remaining monocyte-depleted PBMCs were flushed from the magnetic column, and DNA was isolated using the DNeasy Blood and Tissue Kit (Qiagen, Hilden Germany). RNA was isolated from primary monocytes using the Direct-zol RNA Isolation Kit (Zymo Research, Irvine California), and then DNase treated using the TURBO DNA-free kit (Invitrogen, Carlsbad California). The purity of the isolated monocytes was measured by flow cytometry using the iCyt Synergy SY3200 cell sorter (Sony Biotechnology, Inc, San Jose California), staining with APC/Cy7 anti-CD14 (BioLegend, San Diego California). Monocyte purity was found to be greater than 90%.

### DNA and RNA Sequencing

RNA integrity and concentration was verified using the Agilent Bioanalyzer (RIN> 8) (Agilent Technologies, Santa Clara California). A minimum of 500ng RNA was processed per sample. RNA was ribo-depleted using the NEBNext rRNA Depletion Kit (Human/Mouse/Rat) (New England Biolabs, Ipswich Massechusetts), and sequencing libraries were prepared for every DNA and RNA sample. Sequence-specific magnetic bead capture was performed on DNA and RNA libraries according to the manufacturer’s instructions, using the custom-designed probes (SeqCap EZ Choice XL Library System, Roche Nimblegen, Inc, Madison Wisconsin). Samples were multiplexed, four samples per capture reaction. All post-capture gDNA libraries were sequenced in one lane, while all post-capture cDNA libraries were sequenced in another. All samples were sequenced with the Illumina HiSeq 2500, with paired, 125 bp reads.

### Developing Individualized Genotypes

DNA reads were quality trimmed using Trimmomatic, then aligned to the hg19 reference sequence using the Burrows-Wheeler aligner (BWA MEM) ^25^. Duplicate sequences were then removed using Picard, and indels were processed using GATK ^26–28^. From these aligned reads, SAMtools was used to generate an mpileup file, then VarScan mpileup2snp was used to identify SNPs ^29, 30^. For each individual, SNPs were called based on variation from the reference genome (hg19), and all called SNPs have a total read depth of at least 10 and a maximum variant calling p value of 0.01. For all heterozygous SNPs, each allele also has a minimum variant-supporting read depth of 2, a minimum average variant-supporting read base quality of 20, and a minimum allele frequency of 0.2. From these identified and quality filtered SNPs, individualized lists of variants were created for each sample. On average, 23,575 heterozygous variants were identified in the MHC region per individual.

### RNA Alignment and Assembly

RNA sequencing reads were quality trimmed using Cutadapt, then aligned to the human reference genome (hg19, chromosome 6, RefSeq transcriptome annotation) using GSNAP ^16, 31^. Alternate haplotypes for chromosome 6 in the reference genome were not used for alignment, to prevent errors from multi-mapped reads. RNA reads were aligned in a SNP-tolerant manner, meaning that variants that were called from the DNA sequencing alignment were not included in the mismatch penalty scores for RNA reads. Reads that successfully aligned to the target region were assembled into transcripts using StringTie, guided by the Ensembl transcriptome annotation ^32^.

### Identification of Novel Transcripts

During RNA assembly, transcripts were annotated using an Ensembl reference. To identify novel transcripts, we followed the following workflow (Figure S2). All transcripts that were successfully annotated using the Ensembl reference during alignment were excluded. Using CuffCompare, the remaining transcripts were annotated using Gencode Comprehensive v25 (hg19) as a reference ^33, 34^. All transcripts were assigned class codes based on their relation to transcripts in the reference. All transcripts that were assigned the class codes I (intronic), J (novel isoform), U (intergenic), and X (antisense) were identified as novel, while transcripts containing all other class codes were defined as not novel. The remaining novel transcripts were annotated with CuffCompare using the human lincRNA catalog ^35^. The transcripts that were found to be novel using all three references were next filtered to include only transcripts with an FPKM of 0.1 in two or more samples.

The coding potential of each novel transcript was analyzed using the Coding Potential Analysis Tool (CPAT) ^36^. The sequence of each novel transcript was determined using genomic coordinates determined by StringTie and the sequence of the reference genome, and these sequences were used to determine coding potential for each transcript. Transcripts with a coding probability of 0.364 or greater were defined as putatively coding, while all others were defined as noncoding. All novel transcripts were categorized based on their Gencode annotation class codes and by these coding predictions.

### Predicted Function of Coding Genes

Of the five putative coding transcripts that did not share exons with known genes, structure and function were predicted using IntFOLD3. Using the open reading frame predicted by CPAT, the putative amino acid sequence was determined from the transcript sequences using A plasmid Editor (ApE). Using the IntFOLD3 pipeline, we predicted the tertiary structure of the novel peptides, guided by sequence homology with known proteins ^37^. In addition, putative ligand binding domains and gene ontology term annotation were predicted using the FunFOLD pipeline, which is integrated into IntFOLD3.

### Retroviral Element Sequence Alignment

The five novel putative coding intergenic transcripts described were categorized based on their alignment to retroviral sequences. The predicted open reading frames of each of the five transcripts were translated into a protein sequence. Two transcripts that were isoforms of the same gene shared an open reading frame, so four protein sequences were generated. These sequences were aligned to the human proteome using protein-protein BLAST with the non-redundant protein sequences database^13^. Each of these four sequences aligned to known retroviral elements (E Value < 1 x 10^−10^). Sequence alignments were visualized using MView^38^.

### Allele Specific Expression

Allele specific expression of aligned transcript and genomic reads at each heterozygous SNP was assessed using GATK ASEReadCounter under the default settings, which includes a read downsampling step ^26, 27^. Only alignments with base quality and mapping quality no lower than 20 were used. The read counts for each transcript were then normalized to the allelic variance found in the genomic read counts. Before transcript read counts were evaluated for allele-spcecific expression they were normalized to genomic allele-specific read counts. The DNA allelic imbalance (AI) ratio was first calculated for both the reference and alternate allele of each variant as follows: 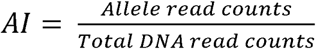. Read counts for both alleles of each variant was then calculated from the following formula:

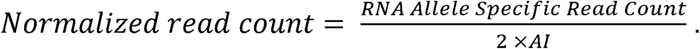

SNPs containing allelic imbalance were defined by a chi-squared test p value less than 0.05, calculated based on the normalized read counts. Relative allelic imbalance for all expressed heterozygous SNPs was calculated as the reference SNP expression: total expression ratio. Heterozygous SNPs and relative reference allele expression for all individuals were merged based on SNP position and reference allele. Allele specific expression at rs76546355 was also confirmed using the program bam-readcount ^39^.

### Co-expression Analysis

Co-expression networks were assigned using the R package WGCNA^12^. This package clusters every sequenced transcript based on the normalized read counts (FPKM) in all twelve samples, using a weighted correlation network analysis. For initial quality filtering, all transcripts that were missing from more than one half of all samples were removed from analysis; 320 transcripts were removed. Samples were then clustered according to transcription patterns to remove any outlier samples; however, no outlier samples were observed. From this filtered set of 2753 genes, a co-expression network was created, with a soft-thresholding power of 7, a dendrogram cut height of 0.25, and a minimum cluster size of 30 transcripts. All transcripts were categorized within an expression dendrogram, then successfully assigned to a co-expression cluster. A total of fourteen clusters were defined. The genomic localization of each cluster was visualized using Circos^40^.

### Transcription Factor Binding Site Enrichment Analysis

Transcription factor binding site enrichment analysis was performed for each of the fourteen co-expression clusters using Genome*Runner* Web^41^, which compares the genomic coordinates of each transcript to the genomic positions of known transcription factor binding sites, using a database that includes the non-cell specific binding patterns of 161 transcription factors, measured via transcription factor ChIP-seq distributed by ENCODE. The coordinates for the promoter region of each gene in each coexpression cluster was used as imput, defined as the 1500bp preceding transcriptional start sites. As a background for enrichment analysis, we included the promoter region of every gene within the MHC region, as annotated by the UCSC known genes list, and also included the novel genes that we have described. The UCSC known gene list contains an aggregation of gene annotations from across the RefSeq, GenBank, CCDS, Rfam, and tRNAscan-SE databases.Transcription factor enrichment was calculated for each co-expression cluster individually, and a cluster was called enriched for a specific transcription factor when an increased frequency of the target was observed in the cluster compared to the background (OR>1, chi-square p<0.05). Nine of the fourteen co-expression clusters were enriched for specific transcription factors.

### HLA Typing

HLA allelotypes for each sample were determined using BWAkit. This pipeline calls types by aligning reads to each HLA gene using the BWA-MEM algorithm, and comparing the exons of each gene to alleles defined by IMGT/HLA. The called types (**Table S4**) are defined as the alleles that have minimal exonic mismatch with the individual’s sequence.

### Sanger Sequencing

Allele-specific expression was validated by Sanger sequencing for the target variant rs76546355. RNA was saved from each individual before sequence-specific capture and was converted into cDNA using the Verso cDNA Synthesis Kit (Thermo Fisher Scientific Inc, Waltham, Massachusetts). This cDNA was then amplified via PCR, using primers that flank the target SNP (forward primer: TGCTTGCCTGTTGTGAGATG, reverse primer: AAGCAACAGTAATTTGGATCTTCC). The proportion of each allele represented in this PCR product was estimated using a Sanger sequencing trace file for each sample.

## Author contributions

E.G. contributed to experimental design, performing the experiments and analyzing the data, and drafting the manuscript; W.W. contributed to performing data analysis, and editing the manuscript; A.H.S. conceived the study, contributed to experimental design, data analysis, and drafting the manuscript.

## Supplementary Material

**Figure S1:**
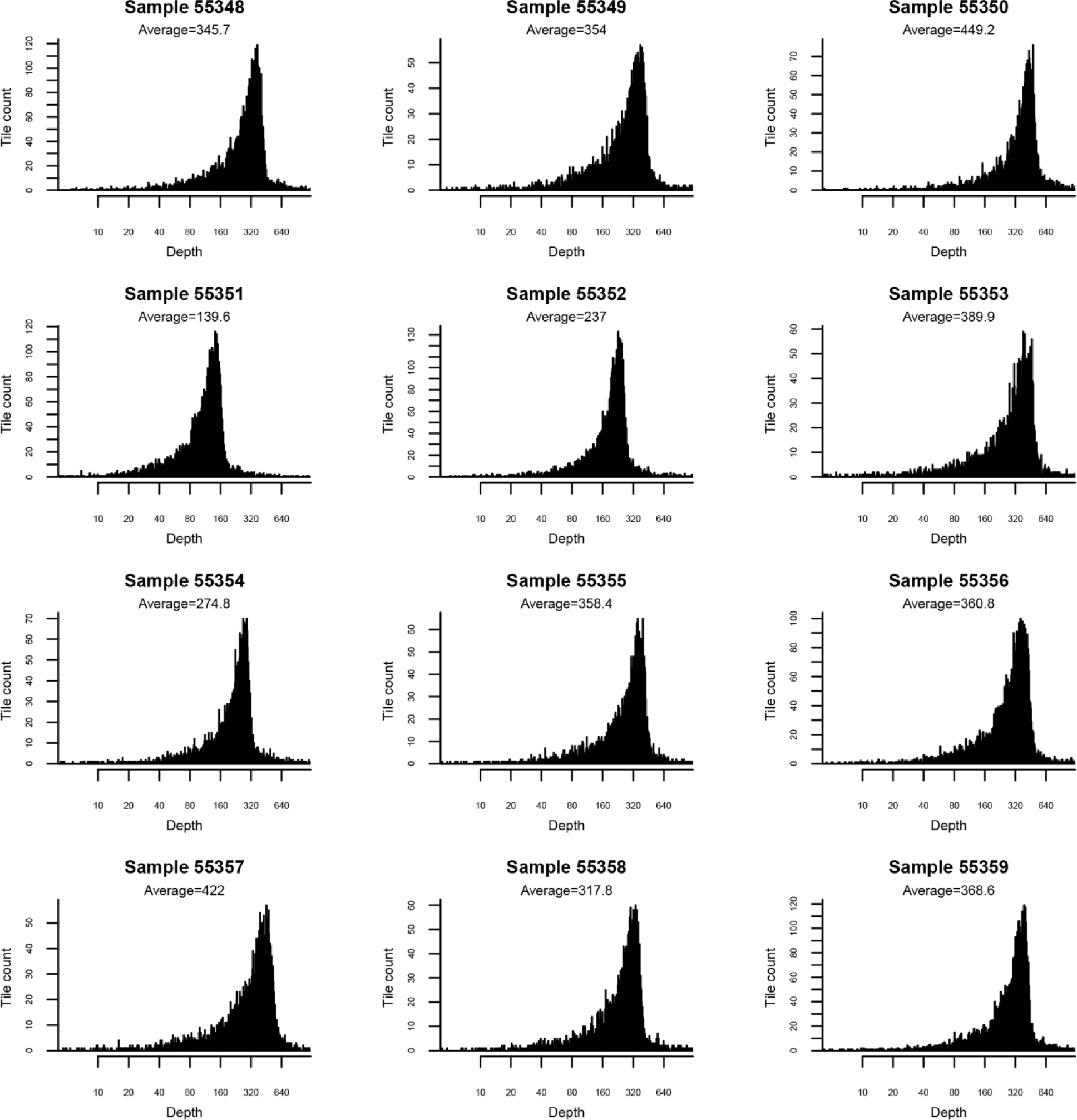
Distribution, mean, and median DNA read depth in tiled regions within the MHC in each sample.

**Figure S2:**
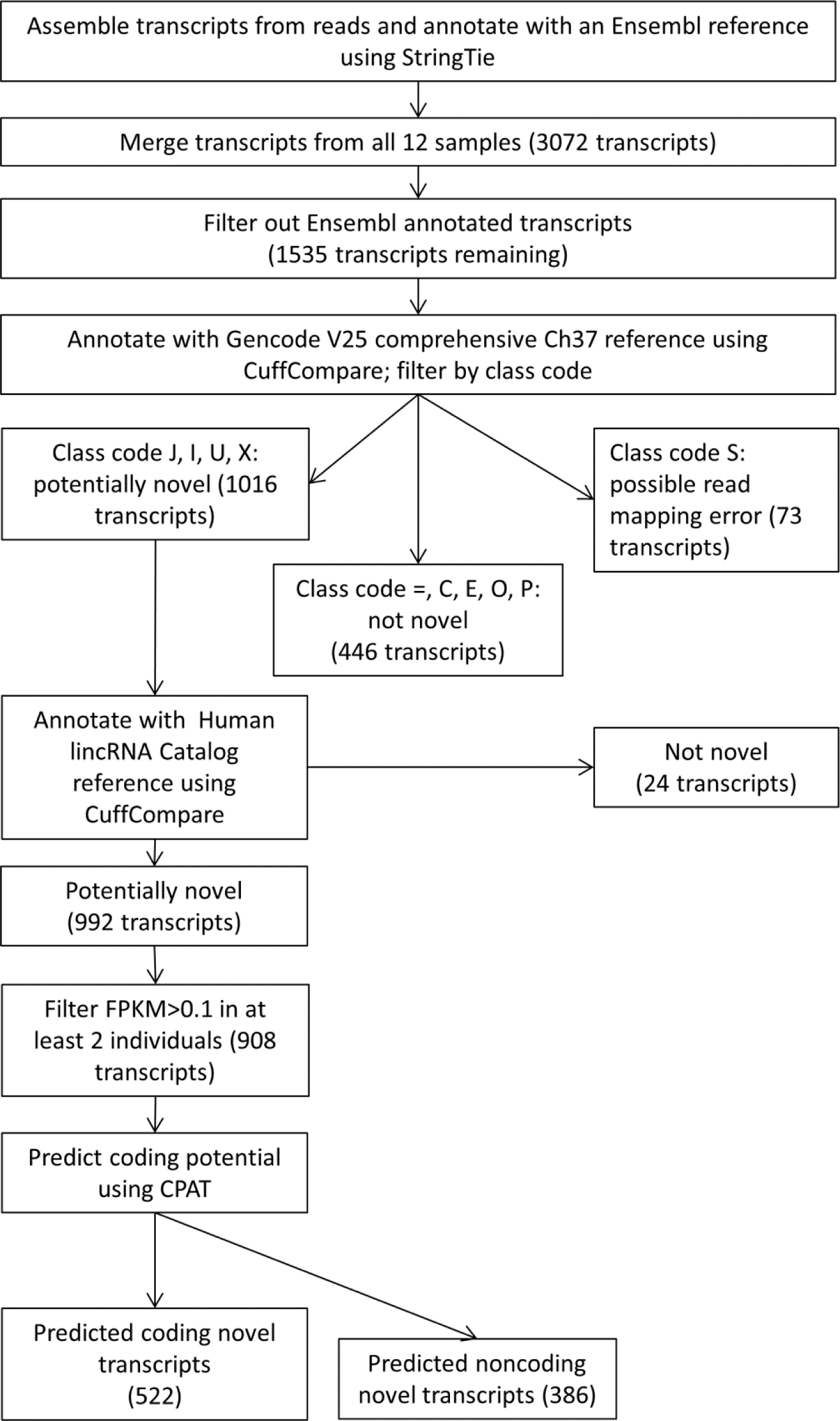
A flow chart depicting the workflow used for identifying novel transcripts from assembled transcripts. In total, 908 novel transcripts were identified. Novel transcripts were not present in the Ensembl, Comprehensive Gencode Release 25, or the Human lincRNA Catalog reference transcriptomes. All novel transcripts were identified in a minimum of two independent samples, and met a relative transcription level threshold (FPKM > 0.1).

**Figure S3:**
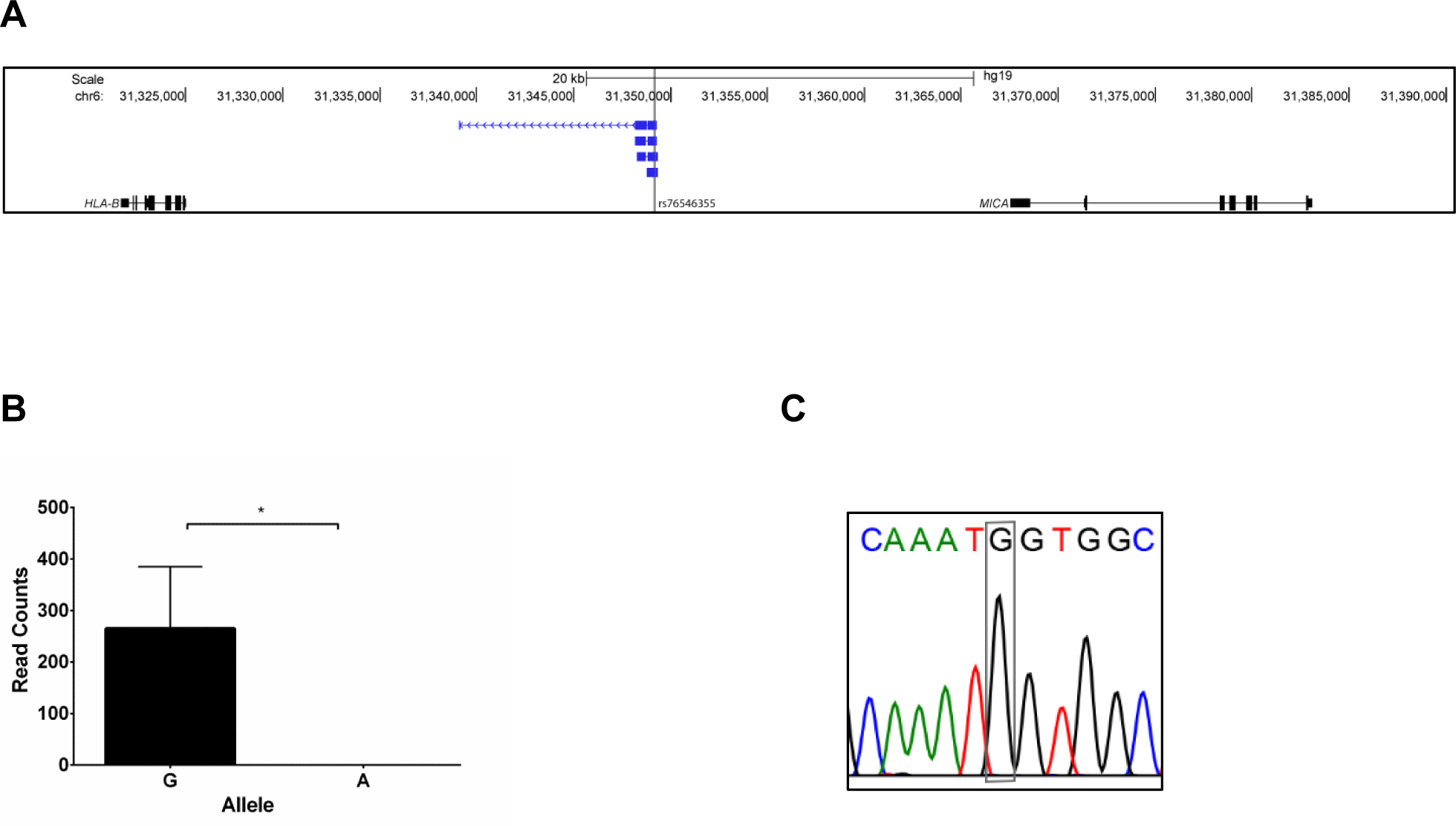
(**A**) The genetic variant rs76546355 (rs116799036) which explains the most robust genetic association for Behçet’s disease and previously thought to be in a non-transcribed genetic region is expressed within four lincRNA transcripts between *HLA-B* and *MICA*. (**B**) RNA sequencing revealed that these lincRNA transcripts are exclusively expressed from the DNA strand with the disease protective allele (allele G), and no expression was detected from the disease risk allele (allele A). RT PCR followed by Sanger sequencing confirmed expression of the novel lincRNA transcripts in this locus, and allele expression imbalance in rs76546355 (a representative chromatogram of seven heterozygous samples is shown) (**C**).

**Figure S4:**
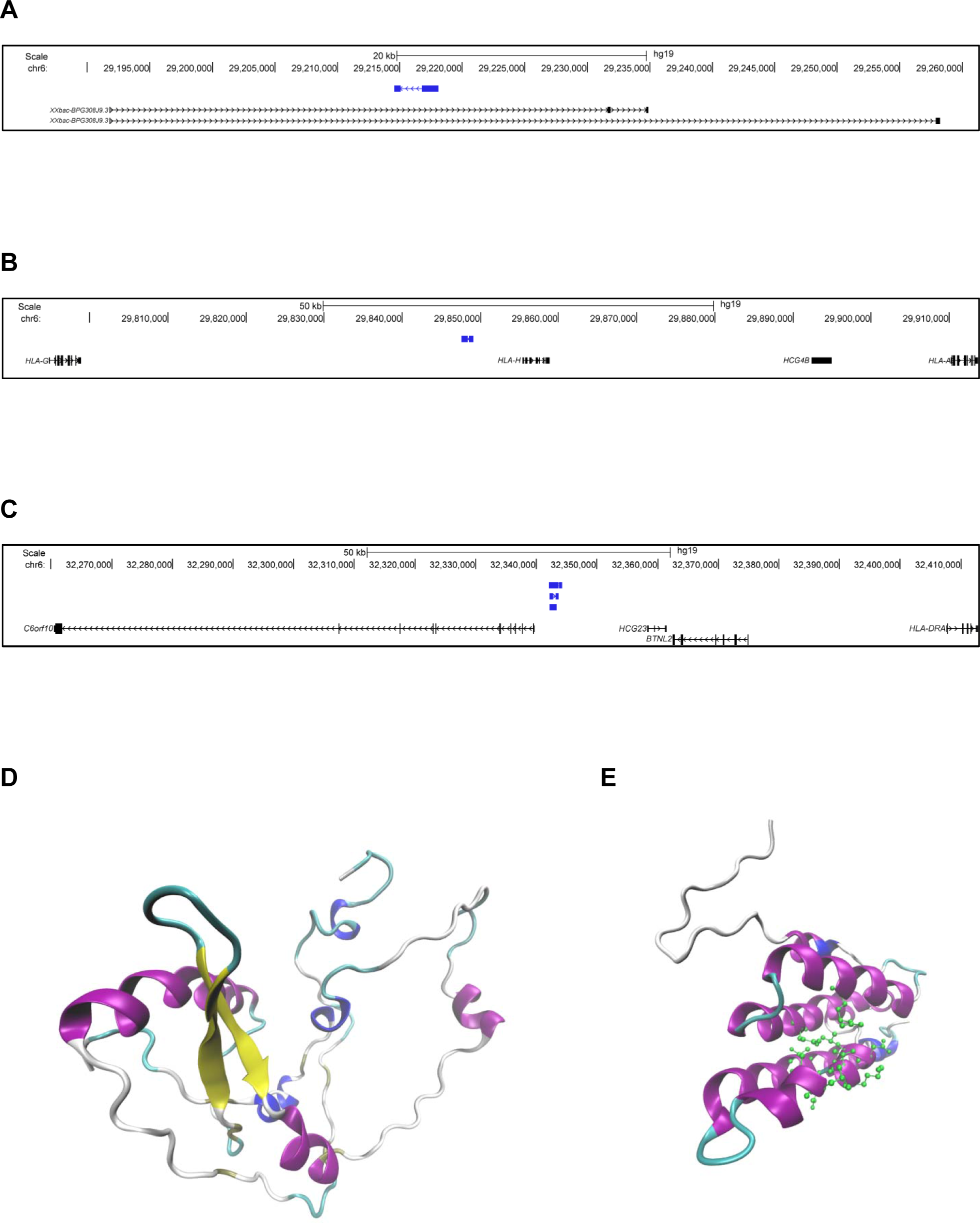
Genomic position (hg19) and predicted protein structure of the three novel protein-coding genes identified in this study. (**A**) One protein-coding novel transcript (blue) contained within the intronic region of the gene *XXbac-BPG308J9.3*. (**B**) and (**C**) depicts novel protein coding transcripts (blue) in intergenic regions, near *HLA-A* and *HLA-DRA*, respectively. Each of these transcripts shares homology with endogenous retroviral elements. (**D**) The Predicted protein structure of transcript A (prediction p value= 1.17 x 10^−4^). This structure shares homology with an endogenous retroviral pol protein, and no predicted ligands are available. (**E**) The predicted protein structure of transcript C (prediction p value= 0.037). This structure shares homology with a retroviral gag protein, and is predicted to bind to a leucine residue (predicted active site amino acids are shown in green). In both **D** and **E** coloring is based on secondary structure: alpha helices are purple, 3-10 helices are blue, beta sheets are yellow, beta bridges are tan, turns are cyan, and coils are white.

**Figure S5:**
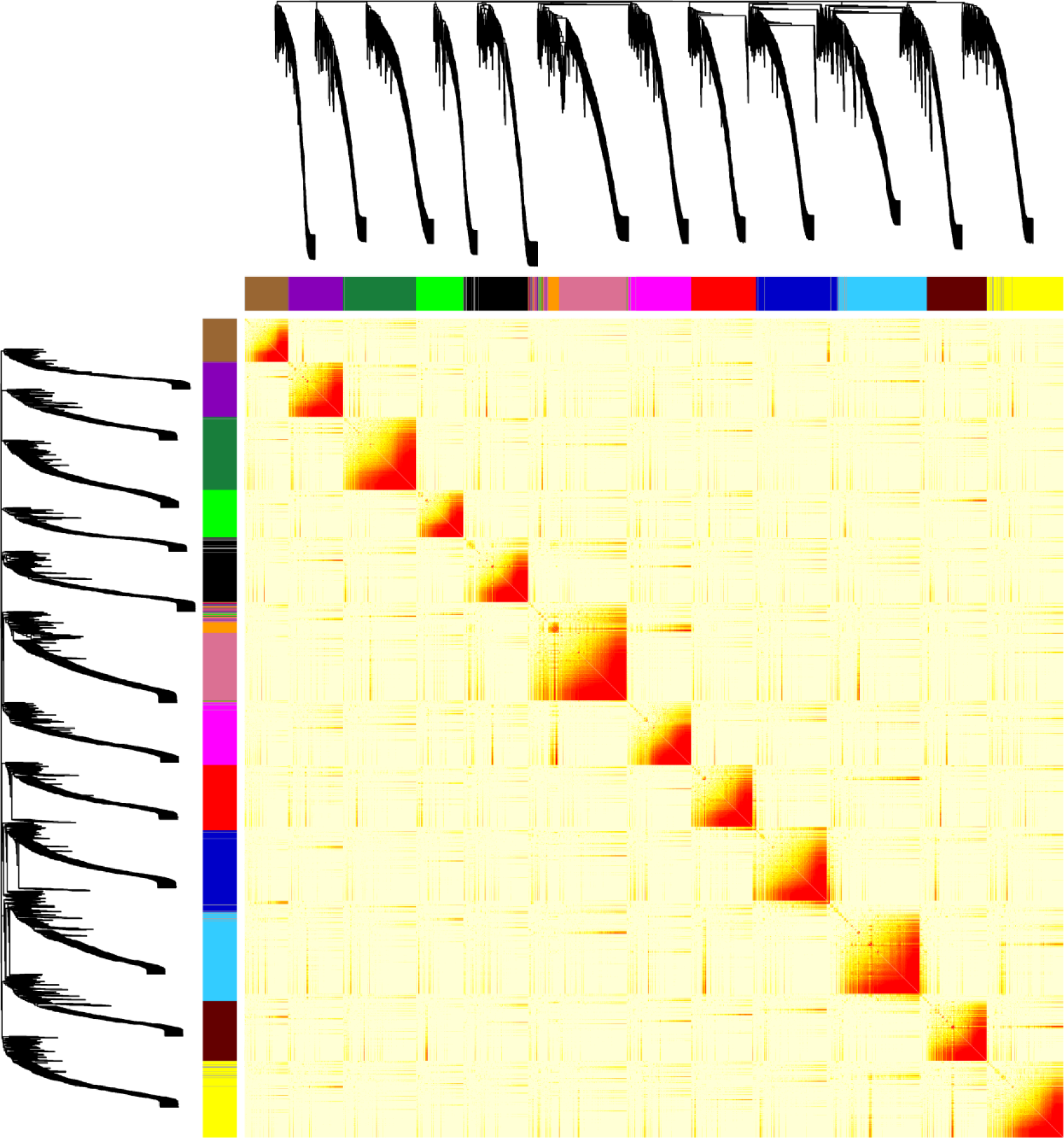
Network heat map depicting expression patterns of all MHC transcripts. Transcripts are aggregated based on co-expression cluster membership, and fourteen discrete colors are used to annotate the clusters on both axes. The precise composition of the co-expression network is seen in the dendrogram before both axes. As demonstrated in this dendrogram, there is a high degree of separation between the clusters in the network. The expression patterns for each transcript are included within the heat map itself (red: high coexpression, yellow: moderate, white: low). In each cluster, transcripts are highly co-expressed, with low or isolated co-expression outside of the assigned cluster.

**Table S1:** A list of all novel transcripts identified within the MHC region in this study, and detected in at least two independent samples.

**Table S2:** The total number of heterozygous SNPs, and the number and fraction of heterozygous SNPs with allelic imbalance in each sample and in all samples in this study.

**Table S3:** Transcribed heterozygous SNPs with evidence of allele specific expression imbalance in at least 2 samples. In each individual separately, heterozygous SNPs were filtered for allele-specific expression based on binomial p value (p<0.05), as calculated by GATK ASEReadCounter. SNPs with allelic imbalance from all samples were then merged based on position, and SNPs with allelic specific expression in two or more samples were included in this table.

**Table S4:** HLA classical alleles in the samples included in this study.

**Table S5:** Multiple sequence alignment of all novel putative coding transcripts with known retroviral element sequences. Protein sequences were translated from the predicted open reading frame of each transcript, and aligned to the human proteome. All alignments (E Value < 1 x 10^−10^) are included.

**Table S6:** Co-expression profile of each aligned gene. Transcripts were grouped into co-expression clusters, differentiated by color assigned. The transcription factor found to be most significantly enriched in each cluster is listed for each transcript.

